# Establishment of a reverse genetic system from a bovine derived Influenza D virus isolate

**DOI:** 10.1101/2021.02.10.430577

**Authors:** Melle Holwerda, Laura Laloli, Manon Wider, Lutz Schönecker, Jens Becker, Mireille Meylan, Ronald Dijkman

## Abstract

The ruminant-associated Influenza D virus (IDV) has a broad host tropism and was shown to have zoonotic potential. To identify and characterize molecular viral determinants influencing the host spectrum of IDV, a reverse genetic system is required. For this, we first performed 5′ and 3′ rapid amplification of cDNA ends (RACE) of all seven genomic segments, followed by assessment of the 5′ and 3′ NCR activity prior to constructing the viral genomic segments of a contemporary Swiss bovine IDV isolate (D/CN286) into the bidirectional pHW-2000 vector. The bidirectional plasmids were transfected in HRT-18G cells followed by viral rescue on the same cell type. Analysis of the segment specific 5′ and 3′ non-coding regions (NCR) highlighted that the terminal 3′ end of all segments harbours an Uracil instead of a Cytosine nucleotide, similar to other Influenza viruses. Subsequent analysis on the functionality of the 5′ and 3′ NCR in a minireplicon assay revealed that these sequences were functional and that the variable sequence length of the 5′ and 3′ NCR influences reporter gene expression. Thereafter, we evaluated the replication efficiency of the reverse genetic clone on conventional cell lines of human, swine and bovine origin by use of an in vitro model recapitulating the natural replicate site of IDV in bovine and swine. This revealed that the reverse genetic clone D/CN286 replicates efficiently in all cell culture models. Combined, these results demonstrate the successful establishment of a reverse genetic system from a contemporary bovine IDV isolate that can be used for future identification and characterization of viral determinants influencing the broad host tropism of IDV.

## 1. Introduction

The *Deltainfluenzavirus* genus is the newest member of the *Orthomyxoviridae* virus family, and harbours the Influenza D virus (IDV) species [1,2]. In 2011, IDV was first identified in a 15-week-old pig suffering from flu-like symptoms such as sneezing and dry cough [3]. Although IDV was initially discovered in swine, subsequent prevalence studies proposed that cattle are the primary host reservoir for IDV with a worldwide distribution [1,4–11]. In addition to cattle and swine, IDV was shown to infect a broad range of other animals, such as feral swine, water buffalo, sheep, horses and camelids [3,8,12–16]. Furthermore, the high seroprevalence among humans with frequent exposure to cattle, as well as the recent demonstration of productive IDV replication in human airway epithelial cells, indicate that the bovine-lineage of IDV has zoonotic potential [17,18].

Reverse genetic systems are an important tool to identify and characterize viral determinants that can influence the viral host tropism [19–23]. The polarity of the segmented viral genome of *Orthomyxoviruses* requires that the 6 – 8 viral genomic segments are individually transcribed in a negative orientation to generate a template for the subsequent production of viral proteins and new progeny virus [24,25]. The first plasmid-based reverse genetic systems for *Orthomyxoviruses* were described in the late 1990s and early 2000s for Influenza A virus [26–28]. The first system consisted of individual plasmids transcribing the negative sense RNA of each viral genomic segment in a RNA polymerase I dependent manner [26]. These plasmids were co-transfected with eukaryotic expression plasmids encoding for the nucleocapsid and RNA dependent RNA polymerase (RdRp) complex proteins to initiate the production of replication competent viruses [26]. This system was later modified by the more efficient bidirectional vector system that incorporates both RNA polymerase I and II dependent transcription of each viral genomic segment and protein, respectively, within the same backbone [27,29]. This bidirectional plasmid-based reverse genetic system has also been described for Influenza B and C viruses [30,31], and more recently also for the swine lineage (D/OK) of IDV [32]. Interestingly, despite cattle being the proposed main reservoir for IDV, no reverse genetic system has so far been established for the more prevalent bovine-lineage [5,33,34].

The aim of this study is to establish a reverse genetic system for a bovine isolate of IDV, which would allow for the molecular characterization of viral determinants influencing the broad host tropism of IDV. For this, we first defined the 5′ and 3′ terminal sequences of the non-coding region (NCR) of the genomic segments from a previously identified Swiss bovine IDV isolate (D/bovine/Switzerland/CN286). This revealed that the first 12 and 11 nucleotides in the 5′ and 3′ terminal NCR regions, respectively, are conserved among all the viral genome segments. However, the sequence length of the NCR genomic segments influences the amplitude of the reporter gene expression in a minireplicon assay. Rescue and subsequent viral replication kinetic experiments with the reverse genetic clone of D/CN286 on conventional human, bovine and swine cell lines, as well as well-differentiated AEC cultures from bovine and porcine origin, revealed robust viral replication. These combined results demonstrate that we have established a reverse genetic system for IDV from a bovine isolate that allows us for performing detailed characterization of viral determinants influencing the host tropism of IDV.

## 2. Materials and methods

### 2.1 Cell culture

The Madin Darby Bovine Kidney (MDBK) cell line was maintained in Eagle’s minimum essential medium (EMEM; (Gibco, Gaithersburg, MD, USA) supplemented with 7% heat-inactivated fetal bovine serum (FBS, Gibco), 2 mmol/L Glutamax (Gibco), 100 μg/mL streptomycin and 100 IU/mL penicillin (Gibco). Human rectal tumour 18G (HRT-18G, American Type Culture Collection, Manassas, VA, USA (ATCC)), Swine testicular (ST, ATCC) cells and human embryonal kidney (HEK) 293-LTV (LTV-100; Cellbiolabs, San Diego, CA, USA) cell lines were maintained in Dulbecco’s minimum essential medium (DMEM; Gibco) supplemented with 5% (HRT-18G) or 10% (ST and 293T-LTV) FBS, 100 μg/mL streptomycin and 100 IU/mL penicillin (Gibco). Well-differentiated airway epithelial cell cultures of bovine and swine origin were established from post-mortem tracheobronchial epithelial tissue and maintained according to the protocol by Gultom *et al.* (2020) [35]. All cell cultures were maintained at 37 °C in a humidified incubator with 5% CO_2_.

### 2.2 Determination of the 5′ and 3′ NCRs by Rapid amplification of cDNA ends (RACE)

The nucleotide sequences of the 5′ and 3′ NCRs of the D/bovine/Switzerland/CN286 (D/CN286) virus isolate were determined by an in-house protocol, based on the 5′ and 3′ RACE kit (2^nd^ generation, Roche, Basel, Switzerland). Viral nucleic acids were extracted from 140 μl of the virus transport medium derived from the nasopharyngeal swab using the viral RNA isolation kit according to the manufacturer’s protocol (Qiagen, Venlo, The Netherlands). For the 3′ RACE, isolated viral RNA was poly-adenylated using *E. coli* poly-A polymerase (New England Biolabs, NEB, Ipswich, MA, USA) for 30 minutes at 37 °C, followed by a reverse transcriptase (RT)-step with Moloney Murine Leukemia Virus RT (MMLV-RT, Promega, Madison, WI, USA) in the presence of RNase inhibitor (RNAsin plus, 10U, Promega) by incubation for 1 hour at 42 °C and 20 minutes of 65 °C using the tagRACE_dT16 oligonucleotide. For 5′ RACE, instead of the tagRACE_dT16 oligonucleotide, a segment-specific reverse oligonucleotide was used as described above, followed by nucleic acid purification and poly-adenylation of the cDNA with terminal transferase (NEB) for 30 minutes at 37 °C followed by 10 minutes heat-inactivation at 70 °C. For both the 3′ and 5′ RACE, an initial touchdown PCR was performed with Hotstart Taq mastermix (Qiagen) with segment-specific primers and tagRACE_dT16 reverse oligonucleotides **(Supplementary Table 1),** using the following cycle profile; 95 °C for 15 min; 15 cycles of 94 °C for 30 s, 65 °C touchdown to 50□Ø°C for 1 min, 72 °C for 1 min; 25 cycles of 94 °C for 30 s, 50 °C for 1 min, 72 °C for 1 min. After the first amplification round, the segment-specific PCR products were 100-fold diluted in nuclease free water and used as template in a nested PCR using segment-specific oligonucleotides, in combination with the tagRACE adapter oligonucleotide, using the touchdown PCR protocol **(Supplementary Table 1).** Thereafter, PCR products were purified using the Nucleospin Gel and PCR clean-up kit (Macherey-Nagel, Oensingen, Switzerland) from which the nucleotide sequence was determined using Sanger sequencing (Microsynth, Balgach, Switzerland). Sequence analysis was performed in Geneious Prime (Biomatters, Auckland, New Zealand, v2020.2.4). The PB2 (5′), HEF (5′), and NS1 (3′) sequences were determined from purified product cloned into the pGEM-T vector by TA-cloning (Promega) according to the manufacturer’s guidelines.

### 2.3 Plasmid construction

#### 2.3.1 Minireplicon reporter constructs

For the generation of minireplicon reporter constructs for each of the seven genomic segments of IDV, the coding sequence of the Gaussia luciferase reporter was amplified from the pMCS-Gaussia (Thermo Fischer Scientific, Darmstadt, Germany) plasmid using a specific primer set **(Supplementary Table 1)**, and CloneAmp HiFi PCR (Takara, Tokyo, Japan) according to manufacturer’s guidelines, using the following cycle profile; 98 °C 10 seconds, 55 °C 10 seconds, 72 °C 10 seconds for 35 cycles. Followed by subcloning the PCR amplicon in Bam-HI (NEB) restriction digested pCW57-tGFP-2A-MCS plasmid, kindly provided by Prof. Dr. Adam Karpf (Addgene plasmid #71783, [36]), using the In-Fusion HD cloning kit (Takara). From the resulting pCW57-GFP-2A-Gaussia plasmid, the tGFP-2A-Gaussia luciferase coding sequence was amplified using oligonucleotides harbouring segment-specific NCR nucleotide sequence overhangs from the IDV D/660 isolate **(Supplementary Table 1)**. The resulting seven different amplicons were individually subcloned into the pHH21 vector, kindly provided by Prof. Dr. Georg Kochs, University of Freiburg, Germany, using the In-Fusion HD cloning kit (Takara). As a transfection control for the minireplicon assay, the tRFP-2A coding sequence was amplified from the pCW57-tRFP-2A-MCS plasmid, kindly provided by Prof. Dr. Adam Karpf (Addgene plasmid #78933, [36]) **(Supplementary Table 1)**, using CloneAmp HiFi PCR (Takara) and was subcloned in the Thymidine kinase Cypridina plasmid (Thermofisher) in-frame with the Cypridina luciferase gene, using the In-Fusion HD cloning kit (Takara), resulting in the pTK-RFP-2A-Cypridina luciferase plasmid. The polymerase complex subunits PB2, PB1, P3 and NP from D/660 and D/CN286 were amplified from the respective bidirectional pHW2000 plasmid constructs using CloneAmp Hifi PCR mix (Takara) with the following cycle profile; 98 °C 10 seconds, 55 °C 10 seconds, 72 °C 20 seconds for 35 cycles followed by subcloning into the eukaryotic expression vector pCAGGs using the In-Fusion HD cloning kit (Takara). All plasmids were midiprepped using the Nucleobond Xtra midi kit (Macherey-Nagel), and verified by Sanger sequencing (Microsynth).

#### 2.3.2 Construction of reverse genetic clone

Two microliters of extracted viral RNA were used as template to amplify the seven genomic segments individually with segment-specific oligonucleotides using the Superscript IV one-step RT-PCR system (Thermo Fisher Scientific), according to manufacturer guidelines. The bidirectional pHW2000-vector, kindly provided by Prof. Dr. Martin Schwemmle, University of Freiburg, Germany, was amplified using CloneAmp HiFi PCR mix (Takara) in a 2-step PCR-program of 98°C for 10 seconds followed by 68°C for 4 minutes, with 35 cycles. Both PCR products were excised from a 1% agarose gel and extracted using the Nucleospin Gel and PCR clean-up kit (Macherey-Nagel) and used as templates for In-Fusion HD cloning (Takara). Positive constructs were identified by colony PCR using the segment-specific PCR primers and Taq green mastermix (Promega) with the following cycle profile; 94°C 5 minutes, followed by 25 cycles of 94°C, 1 minute, 55°C 1 minute, 72°C 2 minutes, and a final elongation step of 7 minutes at 72°C. All plasmids were midiprepped using the Nucleobond Xtra midi kit (Macherey-Nagel), and verified by Sanger sequencing (Microsynth).

### 2.4 Minireplicon assay

One day prior to the minireplicon assay, 293-LTV cells were seeded at a density of 25.000 cells per well, in black-wall, clear bottom 96-well plates (Costar, Corning, NY, USA) in DMEM medium containing 15 mM HEPES (Gibco). For each minireplicon reaction, 50 ng of reporter plasmid containing one of the IDV segment specific NCR was mixed with 100 ng of each polymerase subunit (PB2, PB1 and P3) and 200 ng of NP in the eukaryotic expression vector pCAGGs in 50 μl of Optimem (Gibco), together with 250 ng of pTK-RFP-2A-Cypridina luciferase as transfection control. An equal amount of Optimem containing PEImax (1 mg/ml, Polysciences, Warrington, PA, USA) in a 1:3 DNA:transfection reagent ratio was prepared and added to the overexpression plasmids. After 20 minutes of incubation at room temperature, 20 μl volumes from the transfection mixture were divided into three technical replicates. The polymerase activity based on secreted luciferase activity was monitored, by collecting 50 μl of supernatant and replenishment with fresh medium every 24 hours. Supernatant samples were stored at −80°C for subsequent analysis. In parallel, the polymerase activity was monitored by the detection of fluorescence reporter expression on a Cytation 5 Cell Imaging Multi-Mode Reader (BioTek, Sursee, Switzerland) equipped with a 4x air objective (numerical aperture (NA): 0.13). Four images per well were acquired to cover the entire surface of the well and processed and stitched using the Gen5 Image prime software package (v3.08.01). The number of cells that express turbo green fluorescent protein (tGFP, polymerase activity) or turbo red fluorescent protein (tRFP, transfection control) were determined for each individual well using the Gen5 ImagePrime software package (v3.08.01). Gaussia and Cypridina luciferase activities were measured separately in the collected supernatant samples using the Pierce Gaussia or Cypridina luciferase flash assay kits (Thermo Fischer Scientific), respectively, on a Cytation 5 Cell Imaging Multi-Mode Reader (BioTek). Ten μl of supernatant per reaction were loaded into white, half area 96-well plates (Costar), followed by the injection of 15 μl of substrate and direct measurement of the luminescent signal.

### 2.5 Rescue of IDV from reverse genetic plasmids

One day prior to virus rescue, 1 million HRT-18G cells per well were seeded into 6-well plates (Techno Plastic Products AG (TPP), Trasadingen, Switzerland). For every viral rescue, each viral segment in the bidirectional pHW2000 plasmid was diluted to a concentration of 0.2 μg per reaction, followed by transfection using PEImax (1 mg/ml) in a 1:3 DNA:transfection reagent ratio. After 48 hours of incubation at 37°C in a humidified incubator with 5% CO2, the supernatant was removed, and cells were washed twice with MEM medium. After the final washing, cells were supplemented with infection medium, consisting of MEM with 0.5% bovine serum albumin (Sigma-Aldrich), 15 mM of HEPES (Gibco), 100 μg/ml of penicillin and 100 IU of streptomycin (Gibco), and 0,25 μg/ml of bovine-pancreas isolated trypsin (Sigma-Aldrich). Thereafter, cells were incubated for an additional 72 hours at 37°C. Hereafter, the collected supernatant was diluted 10-fold and inoculated onto a confluent layer of HRT-18G cells seeded in a 6-well cluster plate and incubated for 72 hours at 37°C in a humidified incubator with 5% CO_2_. The final virus stock was collected and titrated by tissue culture infectious dosis 50 (TCID_50_) method, as described previously [18]. Rescued viruses were sequence-verified with Sanger sequencing through the amplification of individual genomic segments, as described above.

### 2.6 Nucleotide homology analysis

Nucleotide sequences of each segment of the prototypic D/bovine/Oklahoma/660/2013 (D/660, accession numbers: KF425659-65) were analysed for their homology to the D/CN286 consensus sequences determined by MINion Nanopore sequencing (Glaus *et al.,* Manuscript in preparation) in Geneious prime (v2020.2.4).

### 2.7 Viral replication kinetics

One day prior to infection, the conventional cell lines HRT-18G and ST were seeded at a density of 250.000 cells per well in a 12-well format (TPP), while 125.000 cells were used for the MDBK cell line. Viral infection was performed as previously described, using an MOI of 0.01 for the HRT-18G and ST cell lines, while the MDBK cell line was infected with an MOI of 0.1 [18]. The well-differentiated AEC cultures of porcine and bovine origin were inoculated with 10.000 TCID_50_ per insert. For the conventional cell lines, the viral replication kinetics were monitored for 72 hours in 24-hour intervals, at which 250 μl of supernatant was collected and replenished with fresh medium. While progeny virus production was monitored for 96 hours for the well-differentiated porcine and bovine AEC cultures, the apical wash was collected every 24 hours as described before [18]. After the last time point, cells were fixed with 4% (V/V) neutral buffered formalin and prepared for immunostaining (Formafix AG, Hittnau, Switzerland).

### 2.8 Immunostaining

Formalin-fixed conventional cell cultures and well-differentiated AEC cultures were immunostained according to a previously described protocol [35]. For the detection of IDV-positive cells, cell cultures were stained with a custom-generated rabbit polyclonal antibody directed against the NP of the prototypic D/bovine/Oklahoma/660/2013 strain (Genscript, Piscataway, NJ, USA) [18]. Alexa Fluor^®^488-labeled donkey anti-Rabbit IgG (H+L) (Jackson Immunoresearch) was applied as secondary antibody. For the porcine and bovine AEC cultures cilia and tight junctions were visualized using an Alexa Fluor^®^ 647-conjugated rabbit anti-β-tubulin IV (9F3, Cell Signaling Technology, Danvers, MA, USA) and Alexa Fluor^®^ 594-conjugated mouse anti-ZO-1 (1A12, Thermo Fisher Scientific). All samples were counterstained using 4’,6-diamidino-2-phenylindole (DAPI, Thermo Fisher Scientific) to visualize nuclei. Images for the HRT-18G, ST, and MDBK cell lines were acquired on the EVOS FL auto 2 fluorescence microscopes (Thermo Fischer Scientific), using a 10x air (NA: 0.3) objective. For the AEC cultures, inserts were mounted on Colorforst Plus microscopy slides (Thermo Fisher Scientific) in Prolong diamond antifade mountant (Thermo Fisher Scientific) and overlaid with 0.17 mm high precision coverslips (Marienfeld, Lauda-Königshofen, Germany). Z-stack images were acquired on a DeltaVision Elite High-Resolution imaging system (GE Healthcare Life Sciences, Chicago, IL, USA) using a step size of 0.2 μm with a 60×/l.42 oil objective. Acquired images were cropped and deconvolved using the integrated softWoRx software package and processed using Imaris (Version 9.4.3, Bitplane AG, Zürich, Switzerland) and FIJI [37].

### 2.9 Data representation

All graphs were generated using GraphPad Prism software (Prism, San Diego, CA, USA, v9.0.0) and final figures were assembled in Adobe Illustrator CS6 (v16.0.0). Brightness and contrast of microscopy picture were minimally adjusted and processed identically to their corresponding control using FIJI (v1.53) [37].

## 3. Results

### 3.1 Determination of the 5′ and 3′ NCRs of the genomic segments from a bovine IDV isolate

IDV has been shown to infect a broad range of animal species including cattle, swine and small ruminants, with cattle as the proposed main host reservoir [1]. However, so far no reverse genetic system has been established for a bovine isolate of IDV, that can be used to characterize viral determinants which influences the broad host tropism of IDV among even-toed ungulates. We previously determined the prevalence of IDV in a Swiss cattle cohort and analysed the phylogenetic relationship of 10 clinical isolates to that of other IDV isolates (Glaus *et al.,* Manuscript in preparation). During this study we did not determine the 5′ and 3′ NCRs of each genomic segment. However, because of the importance of the NCR in viral replication and the previous reported sequence discrepancy among different IDV isolates in public sequence repositories, we decided to determine the 5′ and 3′ NCR sequences of the genomic segments from one of our bovine IDV isolates [38]. Based on the Ct-value and available amount of nasopharyngeal material, this was chosen to be the D/Switzerland/bovine/CN286 isolate (Glaus *et al.,* Manuscript in preparation).

We first performed 3′ RACE for each genomic segment and readily observed that all seven genomic segments have a terminal Uracil nucleotide at the 3′ end of the genome, similar to other Influenza viruses, and found that the first 11 nucleotides (UCGUAUUCGUC) of the 3′ NCR are conserved among all genomic segments **(Figure 1)** [39–42], This finding corroborates the previous results from Ishida and colleagues (2020) and independently highlighted that the terminal nucleotide of the 3′ NCR of genomic segments from several isolates are incorrectly annotated in public sequence repositories, including the prototypic D/Oklahoma/Bovine/660 isolate [38]. The results from the 5′ RACE revealed that the first 12 nucleotides are uniform for all seven genomic segments (AGCAGUAGCAAG) and correspond with the public available sequence data for IDV **(Figure 1).** In line with this observation, the constant terminal universal motif in the 5′ and 3′ NCR in our bovine IDV isolate is followed by a segment-specific variable sequence region that seems to be conserved among different IDV isolates **(Figure 1).**

**Figure 1:**
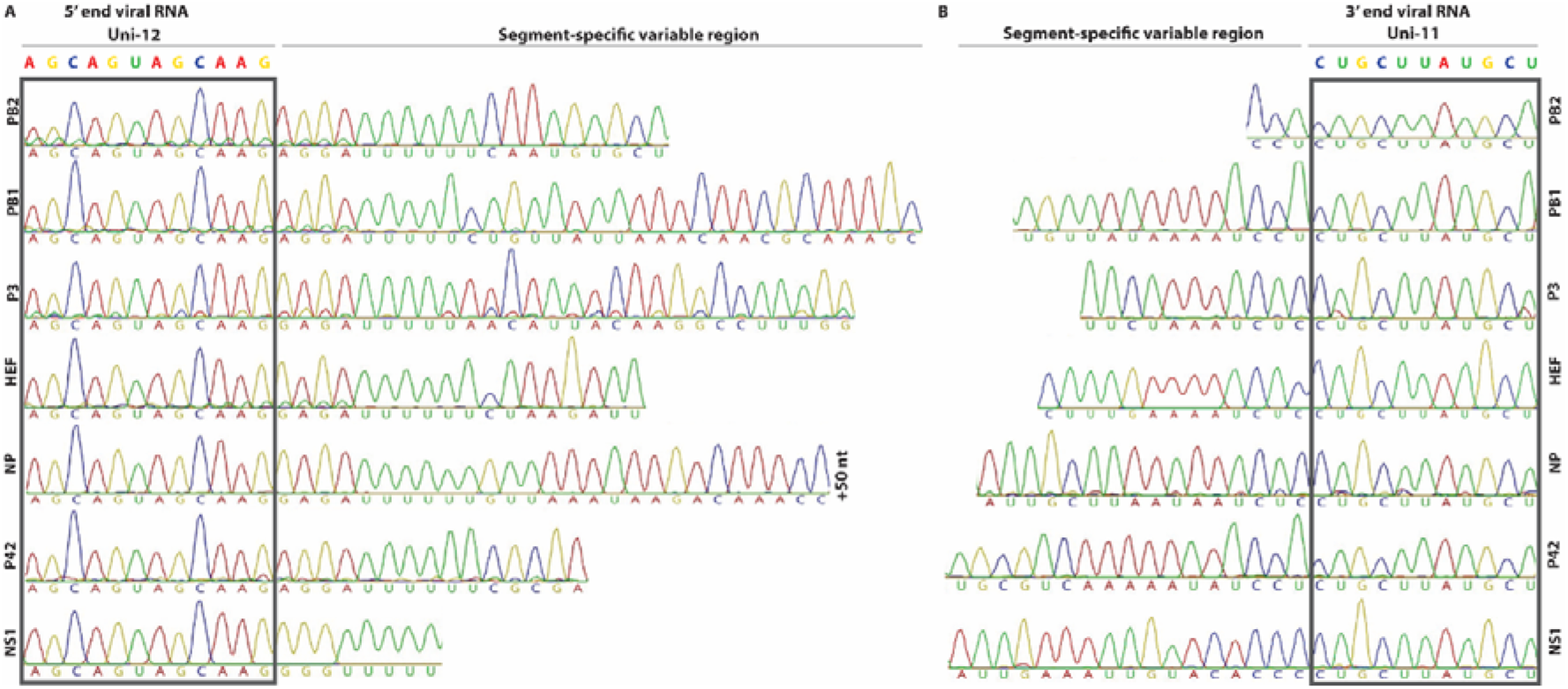
5′ and 3′ RACE sequence results of the non-coding regions from the D/bovine/Switzerland/CN286 virus isolate. The 5′ and 3′ non-coding regions were determined via Sanger sequencing of 5′ and 3′ rapid amplification of cDNA ends (RACE) PCR amplified products. The conserved first 11 nucleotides (UCGUAUUCGUC) and the first 12 nucleotides (AGCAGUAGCAAG) of the 3′ and 5′ NCR, respectively, of the viral RNA are highlighted and adjacent to the segment-specific variable regions.

### 3.2 Analysis of the functionality of the different NCRs by polymerase reconstitution assay

Following the sequence determination of the 5′ and 3′ NCR for each of the seven genomic segments, we wanted to investigate whether the segment-specific NCRs of IDV are functional and whether the nucleotide sequence length influences the transcription and replication efficiency of viral genomic segments. Traditionally, these assays are performed with a plasmid-driven polymerase reconstitution assay (minireplicon). This assay is based on the transfection of eukaryotic expression constructs encoding for the PB2, PB1 and PA/P3 polymerase subunits and the nucleoprotein (NP) in-trans with a plasmid that transcribes a negative strand RNA harbouring a reporter gene (e.g. firefly luciferase) that is flanked by the 3′ and 5′ NCR regions of a viral genomic segment [43,44]. This type of assay is often limited to one timepoint due to the readout method requiring lysis of the transfected cells. To overcome this, we established a reporter construct for IDV that harbours a tGFP-2A-Gaussia luciferase reporter, which allowed us to monitor and measure the influence of the NCR sequence length on gene expression at multiple time points simultaneously via fluorescent- and luciferase-based quantification assays **(Figure 2A).** To control for the transfection efficiency, we established a control plasmid that constitutively expresses a tRFP-2A-Cypridina luciferase and can be used in parallel for the normalization of the expression of both the tGFP and Gaussia luciferase reporter genes **(Figure 2A).**

**Figure 2:**
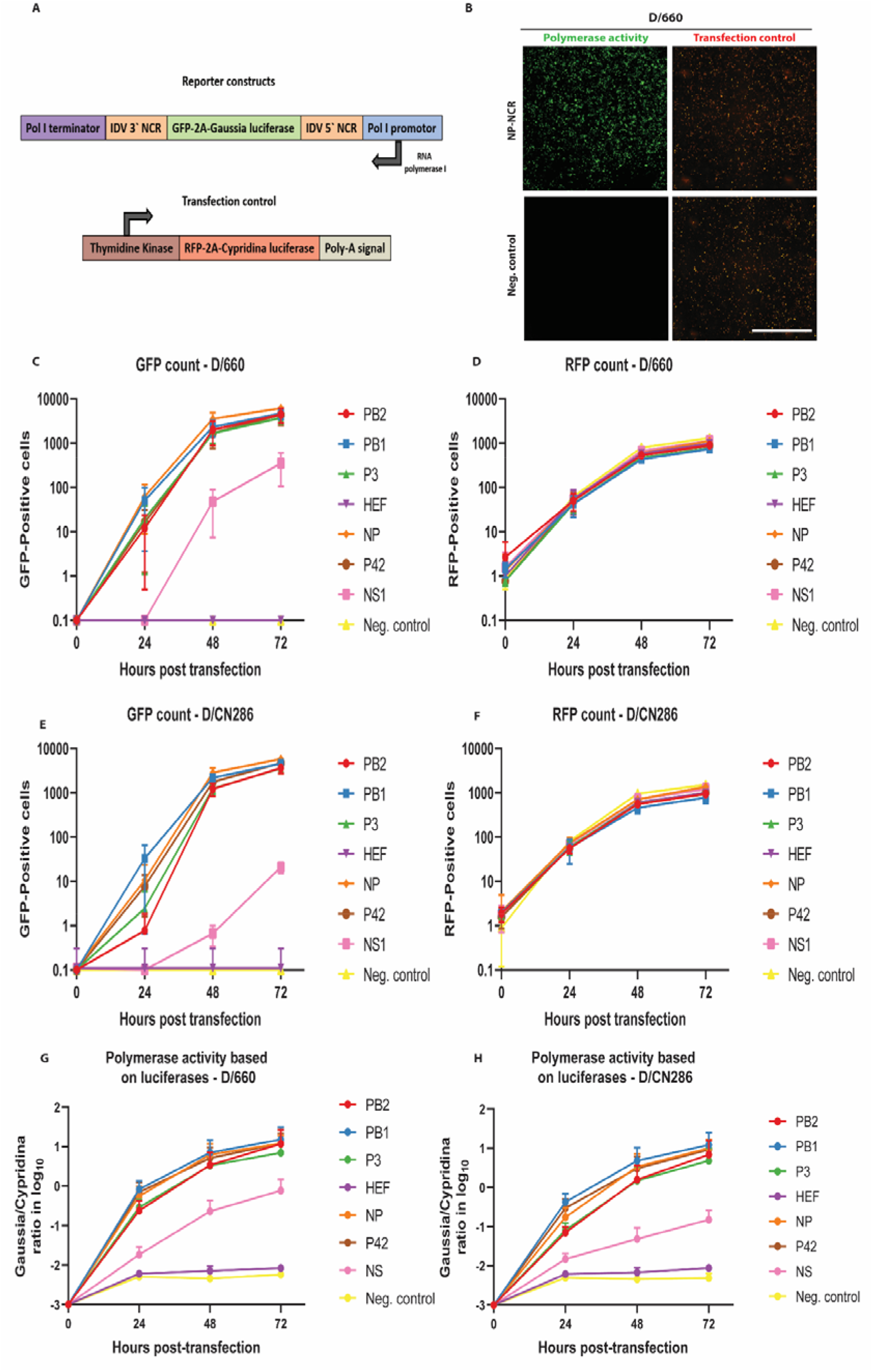
Analysis of the functionality of each segment-specific non-coding region (NCR) of Influenza D virus (IDV) by polymerase reconstitution assay (minireplicon). 293-LTV cells were transfected with expression plasmids encoding for the ribonucleoprotein (RNP) polymerase subunits PB2, PB1, P3 and NP of the D/bovine/Oklahoma/660/2013 (D/660) or D/bovine/Switzerland/CN286 (D/CN286) virus isolates, together with a reporter and transfection control. (A) Illustration of the reporter and transfection plasmids, whereby the tGFP-2A-Gaussia luciferase coding sequence in the reporter plasmid is flanked by the 5′ and 3′ NCR of one specific viral genomic segment. The transfection control plasmid expresses the tRFP-2A-Cypridina luciferase polyprotein under control of the Herpes Simplex virus Thymidine Kinase (TK) promoter. The RNP complex activity was monitored for 72 hours by acquiring images and collecting supernatant samples every 24 hours. (B) Representative microscopy images of the D/660 virus RNP complex activity (green) with the NP NCR reporter and transfection control (red) at 72 hours post transfection. Images are representative of three individual experiments performed in three technical replicates. Scale bar is 2000 μm. The counted GFP **(C, E)** and RFP **(D, F)** positive cells of the respective D/660 **(C, D)** and D/CN286 **(E, F)** RNP complexes. Normalized RNP activity of the D/660 (G) and D/CN286 (H) based on secreted luciferases. Results are displayed as means and SD of three individual experiments performed in three technical replicates.

To demonstrate the functionality of this minireplicon assay, we first monitored the activity of the viral RNP complex of the prototypic D/660 isolates, as this virus replicates in cell cultures. This revealed that the tRFP fluorescent signal could be readily observed 24 hours post-transfection, and that a large number of tGFP positive cells were detected in those cells containing the reporter construct with the NCR of the NP viral genomic segment after 72 hours **(Figure 2B).** Here we observed that the highest reporter activity is detected for the NCR of the NP viral genomic segment followed by those for PB1, PB2, P3 and P42 **(Figure 2C)** with equal transfection efficiencies **(Figure 2D).** In contrast, the reporter activity for NS remained relatively low and the fluorescent signal of the HEF NCR was undetectable, similar to the control cells that lacked a reporter plasmid **(Figure 2C).** These results were corroborated when we analysed the activity of the D/CN286 RNP complex. Despite the fact that several amino acid differ between the D/CN286 and D/660 RNP complex subunits, a relatively similar activity was demonstrated, albeit with a moderately higher activity for the latter **(Figure 2E, F, Supplementary Table 2).** The results obtained with the fluorescent-based assay were corroborated by the luciferase-based assay **(Figure 2G, H).** Interestingly, this revealed that the reporter activity of NS is approximately 10-100-fold lower compared to that of NP, PB1, PB2, PA, and P42, while for HEF this is even lower **(Figure 2G, H).** Because we previously observed that temperature potentially influences the replication kinetics of IDV [18], we also assessed the RNP complex activity with the different NCR reporter constructs at 33°C. We observed that the transfection efficiencies and the number of tGFP positive cells for both the D/660 and D/CN286 RNPs are identical at 33°C in comparison to 37°C **(Supplementary Figure 1)**. However, based on the luciferase activity, we observed that the reporter activity as well as fluorescence intensity were lower at 33°C in comparison to 37°C. This minireplicon assay readout is likely influenced by the temperature dependence of protein folding kinetics **(Supplementary Figure 1)** [45]. Nonetheless, these combined results clearly demonstrate that the NCR of the different constructs are active and that the nucleotide sequence length influences the transcription efficiency of the viral genomic segments.

### 3.3 Establishing a reverse genetic system for IDV

After establishing the functionality of the NCR regions, the individual viral genomic segments from both the bovine D/660 and D/CN286 isolates were cloned into the bidirectional pHW2000-vector for the rescue of a reverse genetic clone of a bovine IDV isolate **(Figure 3A, B)** [27], Sequencing of the obtained pHW2000 plasmids revealed that there were three amino acids between the D/CN286 genetic clone and the consensus sequence from the clinical isolate, namely in the PB1 (D388N), P3 (R350K) and HEF (Q48P). While the plasmids from the D/660 genetic clone were identical to the previous published sequences, except for the terminal Uracil nucleotides in the 3′ NCR.

**Figure 3:**
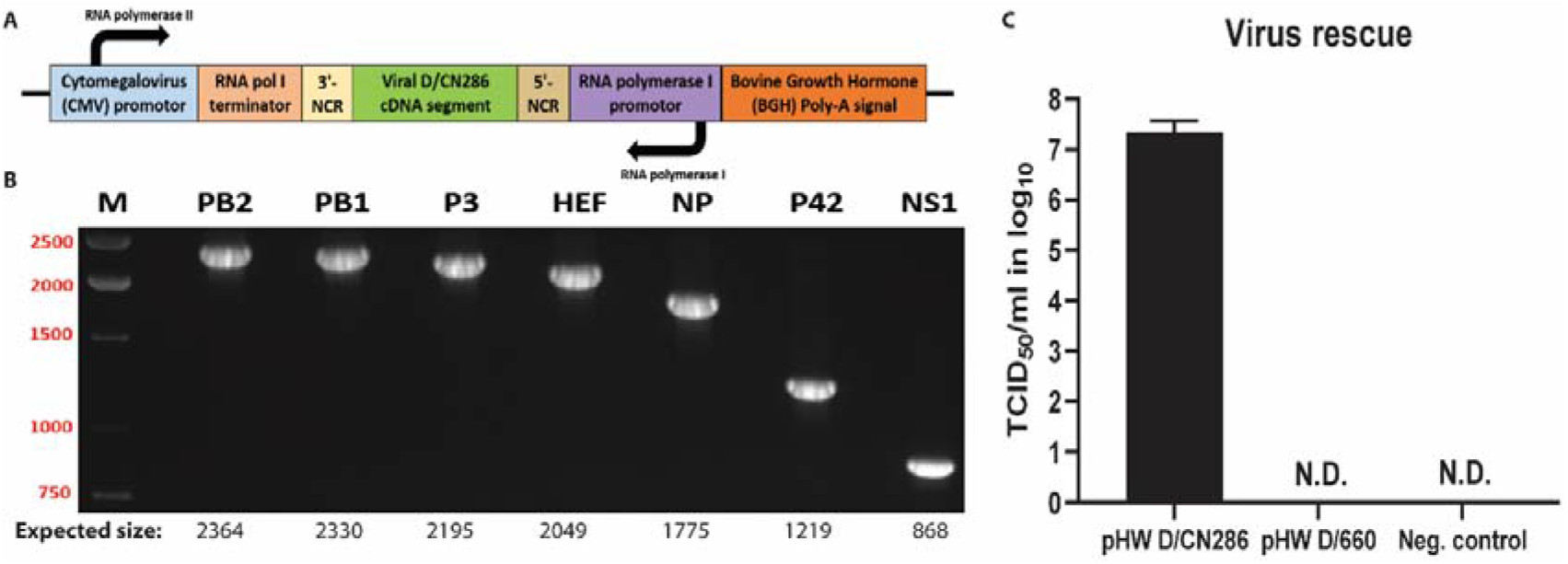
Establishment of a reverse genetic system for IDV. **(A)** A schematic representation of the genetic elements present in the bidirectional pHW2000-backbone. (B) PCR confirmation of cloning the individual viral genomic segments into the pHW2000 plasmid using segment-specific primers. The PCR products were analysed by gel electrophoresis analysis on a 1% agarose gel. The expected fragment sizes are annotated under each lane for each viral genomic segment. (C) Viral titres of the reverse genetic clones D/CN286, D/660, and the negative control after a single passage on HRT-18G-cells. Results are shown as means and SD of three individual experiments performed in two technical replicates per condition. **Abbreviations:** N.D.: not detected.

Following cloning, we transfected the reverse genetic plasmids for rescue of our D/660 and D/CN286 clones into HRT-18G cells, as these cells support the propagation of IDV. After 72 hours of incubation, the supernatant from the transfected cell cultures was collected and passaged upon fresh cultures of HRT-18G followed by viral titration. During the transfection and subsequent passage, we observed no cytopathogenic effect, which was also observed for the cell culture isolate of D/660 (data not shown). However, during the viral titration, that was based on the readout of the agglutination of chicken red blood cells, we measured rescued virus with a viral titre of approximately 10^7^ Tissue culture infectious dose (TCID_50_) per millilitre for the D/CN286 rescued virus, while surprisingly no virus was detected for the D/660 reverse genetic clone **(Figure 3C).** Whole genome sequencing of the passage 1 of the rescued D/CN286 clone revealed that the sequence was identical to that of the reverse genetic plasmids, whereas for D/660 this could not be established.

In the absence of a reverse genetic clone for D/660 and the retrospective identification of another respiratory virus, namely bovine coronavirus, in the CN286 clinical sample, we could not compare the replication kinetics of our rescued D/CN286 reverse genetic clone to the parental strain. Therefore, we instead compared the replication kinetics of the D/CN286 reverse genetic virus with that of the prototypic D/660 cell culture isolate on different cell culture models. We first monitored the viral kinetics on swine (ST), bovine (MDBK) and human (HRT-18G) cell lines for a total duration of 72 hours. This revealed that the replication kinetics of the reverse genetic D/CN286 clone on HRT-18G and ST cell lines is comparable to the cell culture isolate of D/660 **(Figure 4A, B)**. In contrast, the replication kinetics of D/CN286 on the bovine MDBK cell line was approximately 10-fold less efficient compared to the cell culture-adapted D/660 isolate **(Figure 4C)**. In addition to the replication kinetics, we performed immunostaining using a polyclonal NP-directed antibody [18]. This demonstrated that a comparable amount of virus-antigen positive cells could be detected despite the differences in the replication kinetics between both viruses **(Figure 4D-F)**.

**Figure 4:**
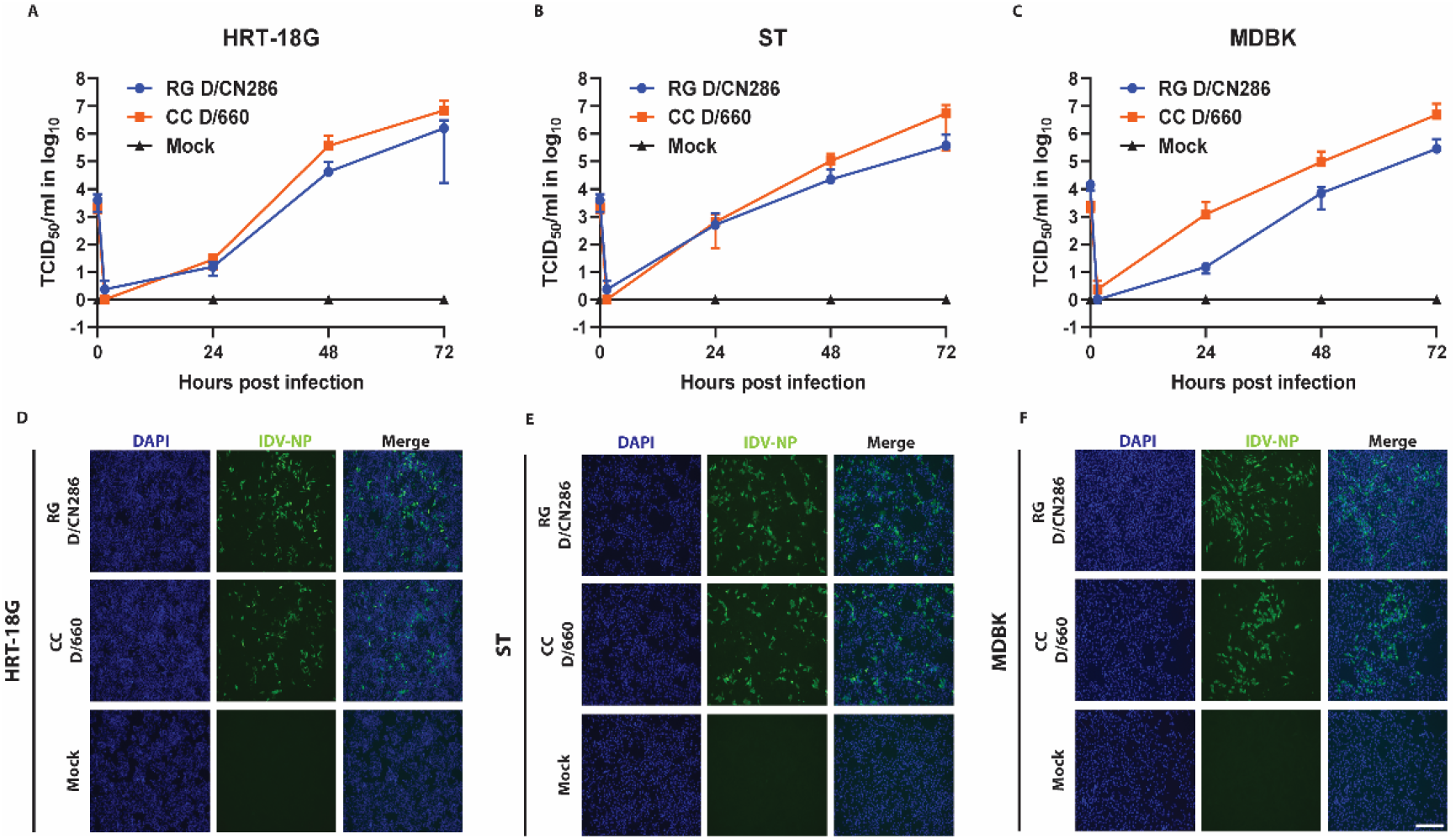
Replication kinetics of IDV on conventional cell lines. The replication kinetics of the reverse genetic D/CN286 clone were compared to the cell culture-adapted D/660 isolate on cell lines of human (HRT-18G, MOI: 0.01), porcine (ST, MOI: 0.01) and bovine (MDBK, MOI: 0.1) origin. Viral kinetics were monitored for 72 hours with 24 hours intervals. Values depicted at 0 hours are the inoculum titres. Viral titres are given as TCID_50_/ml (Y-axis) of the corresponding time points (X-axis) for HRT-18G **(A),** ST **(B)** or MDBK cell lines **(C).** Data points are shown as means and SD of three individual experiments performed in two technical replicates. After 72 hours of viral infection, cells were formalin-fixed and immunostained with the IDV-NP polyclonal antibody (Green) and DAPI (Blue). Representative microscopy images are shown of virus-infected and control culture for HRT-18G (D), ST (E) and MDBK cell lines (F). Images were acquired with a 10x air objective. Scale bar represents 275 μm.

Because the reverse genetic D/CN286 isolate was shown to efficiently replicate in cell lines derived from different species, and we aim to characterize viral determinants that can influence the viral host tropism of IDV, we wondered whether this virus would also infect *in vitro* cultures that recapitulate the natural target site of IDV. To this extend, we inoculated well-differentiated airway epithelial cell (AEC) cultures derived from bovine and porcine origin with 10’000 TCID_50_ and monitored the viral kinetics for 96 hours. This revealed that D/CN286 replicated efficiently in both bovine and porcine AEC cultures, albeit slightly higher in the swine AEC cultures **(Figure 5A, B)**. This was also observed when the swine and bovine AEC cultures were infected with the cell culture-adapted D/660 isolate **(Figure 5A, B)**. Interestingly, this also revealed a similar trend that had been previously observed in the conventional cell lines, namely that the cell culture-adapted D/660 virus replicates more efficiently compared to the D/CN286 reverse genetic clone. However, this effect does not seem to correspond to any difference in cell tropism, as both viruses infected ciliated cells **(Figure 5C, D)**. This observation suggests that potential discrepancies in the replication efficiency are likely associated with the genetic differences between both virus stocks **(Figure 5C, D)**. Nonetheless, these combined results demonstrate that we successfully established a reverse genetic clone from a bovine-derived IDV isolate that efficiently replicates in different cell culture models.

**Figure 5:**
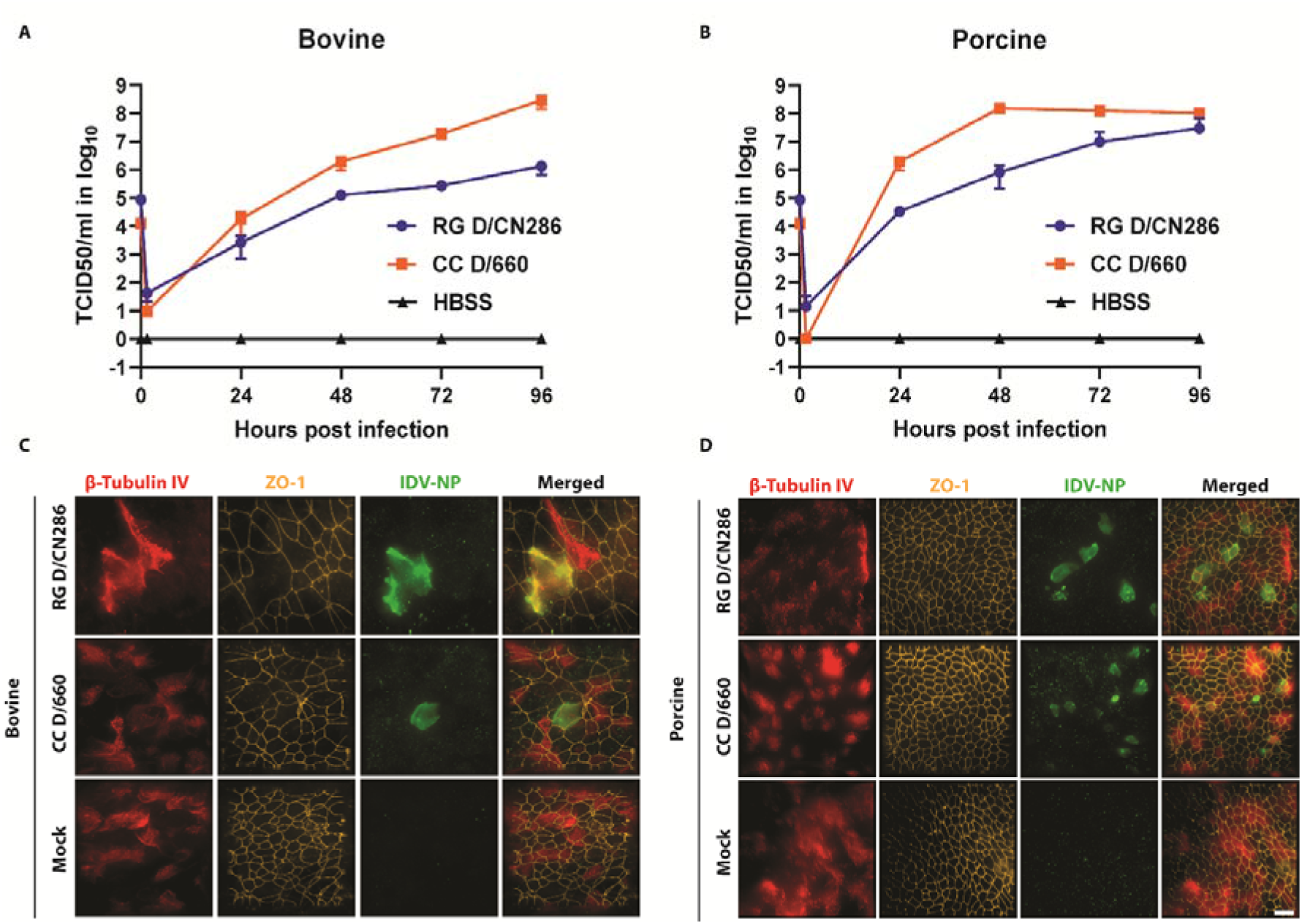
Replication kinetics and determination of the cellular tropism of the rescued Influenza D virus (IDV) clone on well-differentiated airway epithelial cell (AEC) from bovine and porcine origin. AEC cultures were infected with 10.000 TCID_50_ of the rescued IDV clone (RG D/CN286) or cell culture grown IDV (CC D/660). The replication kinetics were monitored for 96 hours of incubation at 37°C and 5% CO_2_, whereby apical washes were collected every 24 hours and analysed by viral titration followed by formalin-fixation and preparation for immunostaining. Viral infectious titres (Y-axis) of the corresponding apical washes are shown as TCID_5_o/ml as indicated hours post infection (X-axis), of the infected bovine (A) or porcine derived AEC cultures (B). Results are shown as means and SD of two technical replicates of one biological donor. Formalin-fixed AEC cultures were immunostained with antibodies visualizing cilia (β-tubulin IV, red), tight junction borders (ZO-1, orange) and virus antigen (IDV-NP, green) of bovine **(C)** and porcine **(D)** derived AEC cultures. Magnification is 60x, scale bar represents 20 μm.

## 4. Discussion

In this study we demonstrate the establishment of a functional reverse genetic system for a contemporary bovine isolate of IDV. Using 5′ and 3′ RACE, we demonstrate that the 11 and 12 terminal nucleotides on the 3′ and 5′ NCRs, respectively are conserved on all viral genomic segments. However, using a novel minireplicon assay, we observed that the sequence length of the conserved segment-specific variable regions influences the RNP driven reporter activity, irrespective of using the RNP complex of the bovine D/660 or D/CN286 virus isolates. Despite the overlapping functionality, viable virus could only be rescued from the pHW-2000 reverse genetic clone of D/CN286, and not for D/660 isolate. The assessment of the replication competency of D/CN286 in different cell culture models, including well-differentiated airway epithelial cultures from bovine and porcine origin, revealed robust virus replication in all models. In combination, these results demonstrate the successful establishment of a reverse genetic system from a contemporary bovine IDV isolate that can be used for future identification and characterization of viral determinants influencing the broad host tropism of IDV.

Previously, two other groups have successfully developed and established a reverse genetic clone for the prototypic swine-derived isolate of IDV (D/OK) [38,46]. However, in line with our results, none of the reports describe the successful establishment of an reverse genetic (RG) clone for the prototypic bovine isolate D/660. Interestingly, Ishida and colleagues and our results independently demonstrate that the terminal nucleotide of the 3′ NCR of the viral genomic RNA contains an Uracil **(Figure 1)** [38], while for the prototypic D/660 isolate the majority of genomic segments are wrongly annotated. However, we demonstrate that the correction of this nucleotide alone does not facilitate the successful rescue of an RG clone of D/660. This indicates that mutations in the coding region of the viral genome are of influence, as we demonstrate that the 5′ and 3′ NCR of the prototypic D/660 isolate are functional. Therefore, it would be of interest to rescue the D/CN286 virus isolate with individual viral genomic segments replaced with one from the D/660. This strategy might reveal which genomic segment negatively influences the rescue of the RG clone of D/660.

In our study, we demonstrate for the first time for IDV that the length of the segment-specific variable 5′ and 3′ NCR sequence has an influence on reporter protein expression. The 5′ and 3′ NCR sequences of the NS and HEF show the lowest reporter gene expression compared to those of the other genomic segments. Interestingly, this phenomenon has also been observed for the phylogenetically related Influenza C Virus, where the sequence length of the variable region in the 5′ NCR of the NS and PB2 segment impairs reporter gene expression [47]. Although we did not perform any mutational analysis on the segment-specific variable 5′ and 3′ NCR in our study, it has been shown previously that a single nucleotide difference can have a detrimental influence [46]. It should however be noted that these results cannot be generalized to the expression levels of viral proteins, as it is well documented for Influenza A virus that both the NCR and gene coding sequence simultaneously can influence transcription efficiency [48,49]. Therefore, further analysis is needed to determine which genomic sequence elements are of importance for IDV transcription.

In our study, we demonstrated that the rescued reverse genetic clone of the bovine D/CN286 isolate replicates in different cell culture models, including well-differentiated airway epithelial cultures from bovine and porcine origin. These results demonstrate, that in addition to our in vitro surrogate model of the human in vivo respiratory epithelium, IDV virus – host interactions can also be studied in analogous models of porcine and bovine origin. However, in comparison to the prototypic bovine D/660 isolate, our RG D/CN286 virus replicated less efficiently on the MDBK cell line and airway epithelial cultures from porcine or bovine origin. This difference might be due to the fact that our RG D/CN286 isolate has 64 amino acid differences compared to the prototypic D/660 isolate, of which 25 are present in the HEF protein **(Supplementary Table 2)**. It is therefore plausible that these differences contribute to the overall lower replication efficiency of our RG clone in our primary bovine and swine AEC cultures. In addition, we previously demonstrated that the cell culture adapted D/660 isolate is composed out of a pool of different genomic variants [18]. Furthermore, the D/660 virus has a greater genomic plasticity to adapt to different host and cellular environments compared to the RG D/CN286 isolate that represents a single genomic variant [50]. Therefore, it would be of interest to study the influence of the 64 amino acid differences in our reverse genetic system and novel minireplicon assay, as well as in well-differentiated airway epithelial cultures from human, bovine and porcine origin. These results might lead to the identification and characterization of viral determinants influencing the broad host tropism of IDV.

In summary, here we show the establishment of a functional reverse genetic system for a contemporary bovine isolate of IDV that will facilitate further characterization of IDV.

## Acknowledgements

We would like to thank Prof. Dr. Georg Kochs and Prof. Dr. Martin Schwemmle from the University of Freiburg, Germany for providing the pHH21- and pHW2000-backbones, respectively.

## References

1. Hause, B.; Collin, E.; Liu, R.; Huang, B.; Sheng, Z.; Lu, W. Characterization of a novel influenza virus strain in cattle and swine: proposal for a new genus in the Orthomyxoviridae family. MBio 2014, 5, 1–10.

2. King, A.M.Q.; Lefkowitz, E.J.; Mushegian, A.R.; Adams, M.J.; Dutilh, B.E.; Gorbalenya, A.E.; Harrach, B.; Harrison, R.L.; Junglen, S.; Knowles, N.J.; et al. Changes to taxonomy and the International Code of Virus Classification and Nomenclature ratified by the International Committee on Taxonomy of Viruses (2018); Springer Vienna, 2018; Vol. 163; ISBN 0070501838471.

3. Hause, B.M.; Ducatez, M.; Collin, E.A.; Ran, Z.; Liu, R.; Sheng, Z.; Armien, A.; Kaplan, B.; Chakravarty, S.; Hoppe, A.D.; et al. Isolation of a Novel Swine Influenza Virus from Oklahoma in 2011 Which Is Distantly Related to Human Influenza C Viruses. PLoS Pathog. 2013, 9, 1–11.

4. Snoeck, C.J.; Oliva, J.; Pauly, M.; Losch, S.; Wildschutz, F.; Muller, C.P.; Hübschen, J.M.; Ducatez, M.F. Influenza D Virus Circulation in Cattle and Swine, Luxembourg, 2012–2016. Emerg. Infect. Dis. 2018, 24, 2012–2013.

5. Ferguson, L.; Eckard, L.; Epperson, W.B.; Long, L.P.; Smith, D.; Huston, C.; Genova, S.; Webby, R.; Wan, X.F. Influenza D virus infection in Mississippi beef cattle. Virology 2015, 486, 28–34.

6. Horimoto, T.; Hiono, T.; Mekata, H.; Odagiri, T.; Lei, Z.; Kobayashi, T.; Norimine, J.; Inoshima, Y.; Hikono, H.; Murakami, K.; et al. Nationwide distribution of bovine influenza D virus infection in Japan. PLoS One 2016, 11, 1–7.

7. Alvarez, I.J.; Fort, M.; Pasucci, J.; Moreno, F.; Gimenez, H.; Näslund, K.; Hägglund, S.; Zohari, S. Seroprevalence of influenza D virus in bulls in Argentina. J. Vet. Diagnostic Investig. 2020, 1–4.

8. Salem, E.; Cook, E.A.J.; Lbacha, H.A.; Oliva, J.; Awoume, F.; Aplogan, G.L.; Hymann, E.C.; Muloi, D.; Deem, S.L.; Alali, S.; et al. Serologic Evidence for Influenza C and D Virus among Ruminants and Camelids, Africa, 1991-2015. Emerg. Infect. Dis. 2017, 23, 2015–2018.

9. Ducatez, M.F.; Pelletier, C.; Meyer, G. Influenza d virus in cattle, France, 2011–2014. Emerg. Infect. Dis. 2015, 21, 368–371.

10. Dane, H.; Duffy, C.; Guelbenzu, M.; Hause, B.; Fee, S.; Forster, F.; McMenamy, M.J.; Lemon, K. Detection of influenza D virus in bovine respiratory disease samples, UK. Transbound. Emerg. Dis. 2019, 66, 2184–2187.

11. Foni, E.; Chiapponi, C.; Baioni, L.; Zanni, I.; Merenda, M.; Rosignoli, C.; Kyriakis, C.S.; Luini, M.V.; Mandola, M.L.; Nigrelli, A.D.; et al. Influenza D in Italy⍰: towards a better understanding of an emerging viral infection in swine. Sci. Rep. 2017, 1–7.

12. Ferguson, L.; Olivier, A.K.; Genova, S.; Epperson, W.B.; Smith, D.R.; Schneider, L.; Barton, K.; McCuan, K.; Webby, R.J.; Wan, X.-F. Pathogenesis of Influenza D virus in Cattle. J. Virol. 2016, 90, 5636–5642.

13. Ferguson, L.; Luo, K.; Olivier, A.K.; Cunningham, F.L.; Blackmon, S.; Hanson-dorr, K.; Sun, H.; Baroch, J.; Lutman, M.W.; Quade, B.; et al. Influenza D Virus Infection in Feral Swine Populations, United States. Emerg. Infect. Dis. 2018, 24, 1020–1028.

14. Zhai, S.; Zhang, H.; Chen, S.; Zhou, X.; Lin, T.; Liu, R.; Lv, D.; Wen, X.; Wei, W.; Wang, D.; et al. Influenza D Virus in Animal Species in Guangdong province, Southern China. Emerg. Infect. Dis. 2017, 23, 1392–1396.

15. Oliva, J.; Eichenbaum, A.; Belin, J.; Gaudino, M.; Guillotin, J.; Alzieu, J.P.; Nicollet, P.; Brugidou, R.; Gueneau, E.; Michel, E.; et al. Serological evidence of influenza D virus circulation among cattle and small ruminants in france. Viruses 2019, 11.

16. Quast, M.; Sreenivasan, C.; Sexton, G.; Nedland, H.; Singrey, A.; Fawcett, L.; Miller, G.; Lauer, D.; Voss, S.; Pollock, S.; et al. Serological evidence for the presence of influenza D virus in small ruminants. Vet. Microbiol. 2015, 180, 281–285.

17. White, S.K.; Ma, W.; McDaniel, C.J.; Gray, G.C.; Lednicky, J.A. Serologic evidence of exposure to influenza D virus among persons with occupational contact with cattle. J. Clin. Virol. 2016, 81, 31–33.

18. Holwerda, M.; Kelly, J.; Laloli, L.; Stürmer, I.; Portmann, J.; Stalder, H.; Dijkman, R. Determining the Replication Kinetics and Cellular Tropism of Influenza D Virus on Primary Well-Differentiated Human Airway Epithelial Cells. Viruses 2019, 11.

19. Herfst, S.; Schrauwen, E.J.A.; Linster, M.; Chutinimitkul, S.; De, E.; Munster, V.J.; Sorrell, E.M.; Bestebroer, T.M.; Burke, D.F.; Derek, J.; et al. Airborne Transmission of Influenza A/H5N1 Virus Between Ferrets. Science (80-.). 2012, 336, 1534–1541.

20. Long, J.S.; Giotis, E.S.; Moncorgé, O.; Frise, R.; Mistry, B.; James, J.; Morisson, M.; Iqbal, M.; Vignal, A.; Skinner, M.A.; et al. Species difference in ANP32A underlies influenza A virus polymerase host restriction. Nature 2016, 529, 101–104.

21. Moreira, É.A.; Locher, S.; Kolesnikova, L.; Bolte, H.; Aydillo, T.; García-Sastre, A.; Schwemmle, M.; Zimmer, G. Synthetically derived bat influenza A-like viruses reveal a cell type-but not species-specific tropism. Proc. Natl. Acad. Sci. 2016, 201608821.

22. Imai, M.; Watanabe, T.; Hatta, M.; Das, S.C.; Ozawa, M.; Shinya, K.; Zhong, G.; Hanson, A.; Katsura, H.; Watanabe, S.; et al. Experimental adaptation of an influenza H5 HA confers respiratory droplet transmission to a reassortant H5 HA/H1N1 virus in ferrets. Nature 2012, 486, 420–428.

23. Neumann, G. Influenza Reverse Genetics—Historical Perspective. Cold Spring Harb. Perspect. Med. 2020, a038547.

24. Dou, D.; Revol, R.; Östbye, H.; Wang, H.; Daniels, R. Influenza A virus cell entry, replication, virion assembly and movement. Front. Immunol. 2018, 9, 1–17.

25. Payne, S. Family Orthomyxoviridae. In Viruses: From Understanding to Investigation; 2017; pp. 197–208 ISBN 9780128031094.

26. Neumann, G.; Watanabe, T.; Ito, H.; Watanabe, S.; Goto, H.; Gao, P.; Hughes, M.; Perez, D.R.; Donis, R.; Hoffmann, E.; et al. Generation of influenza A viruses entirely from cloned cDNAs. Proc. Natl. Acad. Sci. 1999, 96, 9345–9350.

27. Hoffmann, E.; Neumann, G.; Kawaoka, Y.; Hobom, G.; Webster, R.G. A DNA transfection system for generation of influenza A virus from eight plasmids. Proc. Natl. Acad. Sci. 2000, 97, 6108–6113.

28. Fodor, E.; Devenish, L.; Engelhardt, O.G.; Palese, P.; Brownlee, G.G.; García-Sastre, A. Rescue of Influenza A Virus from Recombinant DNA. J. Virol. 1999, 73, 9679–9682.

29. Quinlivan, M.; Zamarin, D.; García-Sastre, A.; Cullinane, A.; Chambers, T.; Palese, P. Attenuation of Equine Influenza Viruses through Truncations of the NS1 Protein. J. Virol. 2005, 79, 8431–8439.

30. Hoffmann, E.; Mahmood, K.; Yang, C.F.; Webster, R.G.; Greenberg, H.B.; Kemble, G. Rescue of influenza B virus from eight plasmids. Proc. Natl. Acad. Sci. U. S. A. 2002, 99, 11411–11416.

31. Pachler, K.; Mayr, J.; Vlasak, R. A seven plasmid-based system for the rescue of influenza C virus. Mol. Genet. Med. 2010, 4, 239–246.

32. Broszeit, F.; Tzarum, N.; Zhu, X.; Nemanichvili, N.; Eggink, D.; Leenders, T.; Li, Z.; Liu, L.; Wolfert, M.A.; Papanikolaou, A.; et al. N-Glycolylneuraminic Acid as a Receptor for Influenza A Viruses. Cell Rep. 2019, 27, 3284–3294.e6.

33. Luo, J.; Ferguson, L.; Smith, D.R.; Woolums, A.R.; Epperson, W.B.; Wan, X.F. Serological evidence for high prevalence of Influenza D Viruses in Cattle, Nebraska, United States, 2003–2004. Virology 2017, 501, 88–91.

34. Collin, E.A.; Sheng, Z.; Lang, Y.; Ma, W.; Hause, B.M.; Li, F. Cocirculation of Two Distinct Genetic and Antigenic Lineages of Proposed Influenza D Virus in Cattle. J. Virol. 2015, 89, 1036–1042.

35. Gultom, M.; Laloli, L.; Dijkman, R. Well-Differentiated Primary Mammalian Airway Epithelial Cell Cultures. In Coronaviruses: Methods and Protocols; Maier, H.J., Bickerton, E., Eds.; Springer US: New York, NY, 2020; pp. 119–134 ISBN 978-1-0716-0900-2.

36. Barger, C.J.; Branick, C.; Chee, L.; Karpf, A.R. Pan-cancer analyses reveal genomic features of FOXM1 overexpression in cancer. Cancers (Basel). 2019, 11.

37. Schindelin, J.; Arganda-Carreras, I.; Frise, E.; Kaynig, V.; Longair, M.; Pietzsch, T.; Preibisch, S.; Rueden, C.; Saalfeld, S.; Schmid, B.; et al. Fiji: An open-source platform for biological-image analysis. Nat. Methods 2012.

38. Ishida, H.; Murakami, S.; Kamiki, H.; Matsugo, H.; Takenaka-uema, A.; Horimoto, T. Establishment of a Reverse Genetics System for Influenza D Virus. J. Virol. 2020, 94, 1–13.

39. Benkaroun, J.; Robertson, G.J.; Whitney, H.; Lang, A.S. Analysis of the variability in the non-coding regions of influenza a viruses. Vet. Sci. 2018, 5.

40. Crescenzo-Chaigne, B.; van der Werf, S. Rescue of influenza C virus from recombinant DNA. J. Virol. 2007, 81, 11282–11289.

41. Zhou, B.; Lin, X.; Wang, W.; Halpin, R.A.; Bera, J.; Stockwell, T.B.; Barr, I.G.; Wentworth, D.E. Universal influenza B virus genomic amplification facilitates sequencing, diagnostics, and reverse genetics. J. Clin. Microbiol. 2014, 52, 1330–1337.

42. Zhou, B.; Donnelly, M.E.; Scholes, D.T.; St. George, K.; Hatta, M.; Kawaoka, Y.; Wentworth, D.E. Single-Reaction Genomic Amplification Accelerates Sequencing and Vaccine Production for Classical and Swine Origin Human Influenza A Viruses. J. Virol. 2009, 83, 10309–10313.

43. Mänz, B.; Dornfeld, D.; Götz, V.; Zell, R.; Zimmermann, P.; Haller, O.; Kochs, G.; Schwemmle, M. Pandemic Influenza A Viruses Escape from Restriction by Human MxA through Adaptive Mutations in the Nucleoprotein. PLoS Pathog. 2013, 9.

44. Welkers, M.R.A.; Pawestri, H.A.; Fonville, J.M.; Sampurno, O.D.; Pater, M.; Holwerda, M.; Han, A.X.; Russell, C.A.; Jeeninga, R.E.; Setiawaty, V.; et al. Genetic diversity and host adaptation of avian H5N1 influenza viruses during human infection. Emerg. Microbes Infect. 2019, 8, 262–271.

45. Guo, M.; Xu, Y.; Gruebele, M. Temperature dependence of protein folding kinetics in living cells. Proc. Natl. Acad. Sci. U. S. A. 2012, 109, 17863–17867.

46. Yu, J.; Liu, R.; Zhou, B.; Chou, T.; Ghedin, E.; Sheng, Z.; Gao, R.; Zhai, S.; Wang, D.; Li, F. Development and Characterization of a Reverse-Genetics System for Influenza D Virus. J. Virol. 2019, 93, 1–14.

47. Crescenzo-Chaigne, B.; Barbezange, C.; Van Der Werf, S. Non coding extremities of the seven influenza virus type C vRNA segments: Effect on transcription and replication by the type C and type a polymerase complexes. Virol. J. 2008, 5, 1–11.

48. Maeda, Y.; Goto, H.; Horimoto, T.; Takada, A.; Kawaoka, Y. Biological significance of the U residue at the −3 position of the mRNA sequences of influenza A viral segments PB1 and NA. Virus Res. 2004, 100, 153–157.

49. Parvin, J.D.; Palese, P.; Honda, A.; Ishihama, A.; Krystal, M. Promoter analysis of influenza virus RNA polymerase. J. Virol. 1989, 63, 5142–5152.

50. Stertz, S.; Palese, P. Genome Plasticity of Influenza Viruses. Genome Plast. Infect. Dis. 2014, 2, 162–177.

