## Supplementary material for "Establishment of a reverse genetic system from a bovine derived Influenza D virus isolate": Figure S1.docx

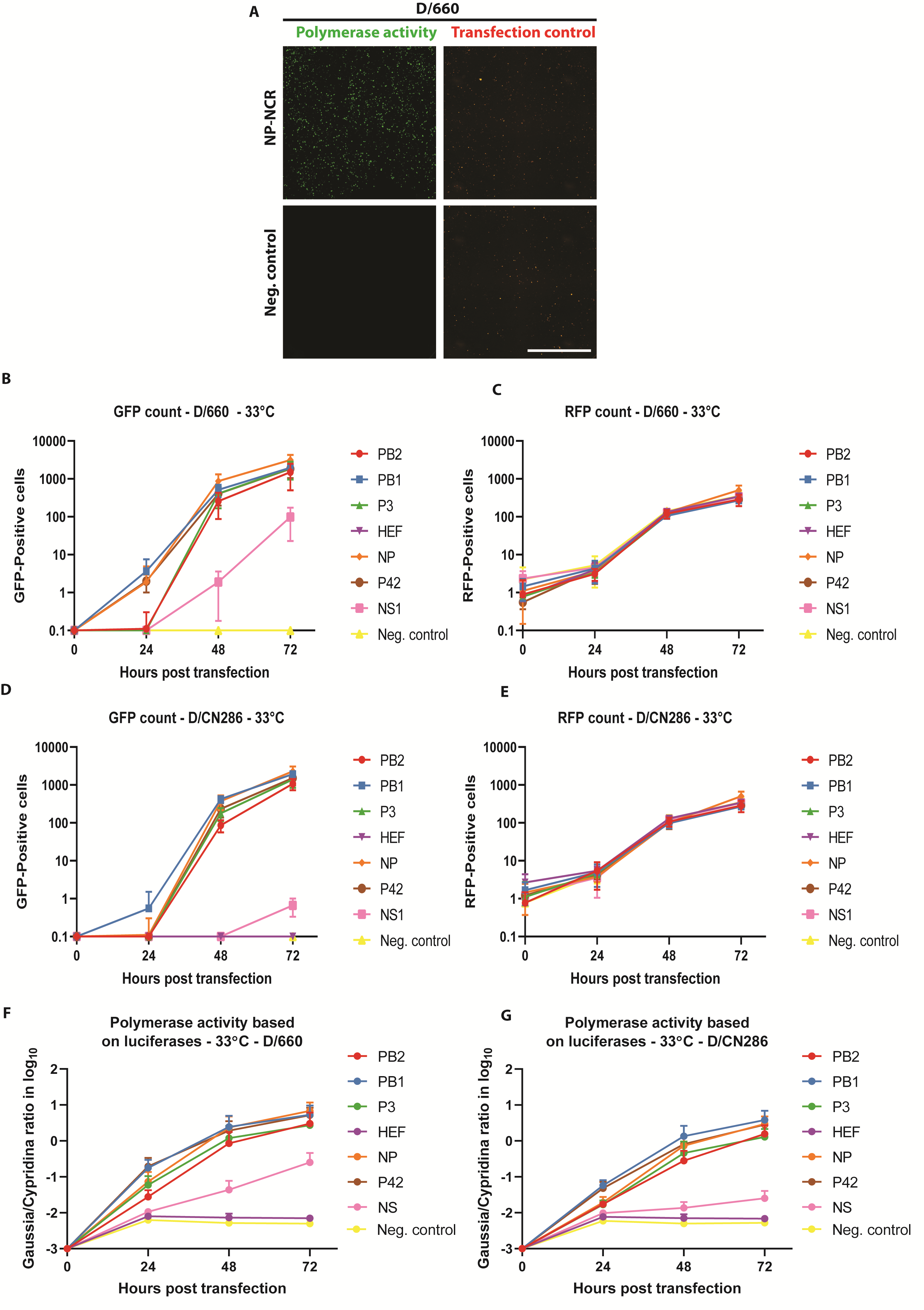

**Supplementary figure 1: Analysis of the functionality of each segment-specific non-coding region (NCR) of Influenza D virus (IDV) by polymerase reconstitution assay (minireplicon) at 33°C.** 293-LTV cells were transfected with expression plasmids encoding for the ribonucleoprotein (RNP) polymerase subunits PB2, PB1, P3 and NP of the D/bovine/Oklahoma/660/2013 (D/660) or D/bovine/Switzerland/CN286 (D/CN286) virus isolates, together with a reporter and transfection control. The RNP complex activity was monitored for 72 hours by acquiring images and collecting supernatant samples every 24 hours.  **(A)** Representative microscopy images of the D/660 virus RNP complex activity (green) with the NP NCR reporter and transfection control (red) at 72 hours post transfection. Images are representative of three individual experiments performed in three technical replicates. Scale bar is 2000 μm. The counted GFP **(B, D)** and RFP **(C, E)** positive cells of the respective D/660 **(B, C)** and D/CN286 **(D, E)** RNP complexes. Normalized RNP activity of the D/660 **(F)** and D/CN286 **(G)** based on secreted luciferases. Results are displayed as means and SD of three individual experiments performed in three technical replicates.
