## Supplementary material for "Establishment of a reverse genetic system from a bovine derived Influenza D virus isolate": Table S1.docx

**Supplementary table 1: Sequence of the used oligonucleotides for cloning constructs.**

Oligonucleotide sequences to determine the nucleotide sequences of the non-coding regions of each specific viral genomic segment by rapid amplification of cDNA ends (RACE):

| **Name** | **Primer sequence (From 5' to 3')** |
| --- | --- |
| **tagRACE_dT16** | GAC CAC GCG TAT CGA TGT CGA CTT TTT TTT TTT TTT TTV |
| **tagRACE** | GAC CAC GCG TAT CGA TGT CGA C |
| **5 prime RACE SSRP PB2** | GAA CTT GCT TCC CAT TGC CCA TC |
| **5 prime RACE SSRP PB1** | CTT CTC TCT ATT GGG AAT ATT CCC G |
| **5 prime RACE SSRP P3** | CCA AAA GAA CGT ATC TTG GGG |
| **5 prime RACE SSRP HEF** | CCT TTT GGT TGA TGA CAC TAG C |
| **5 prime RACE SSRP NP** | CAG AAA ACT TTC CCT GCA TTT GC |
| **5 prime RACE SSRP P42** | TCT CAG GCA TAT GTG CAC CA |
| **5 prime RACE SSRP NS1** | GGC AGC AAT AGT AGT GAA TCC |
| **5 prime RACE PB2 nested** | GAA GGG ATG GGT TTT TCT CTT TTC |
| **5 prime RACE PB1 nested** | CTG TAG GAG TAA GTT CTT GAC AC |
| **5 prime RACE P3 nested** | CAT CAC TTA TAA GGC AGC AAG C |
| **5 prime RACE HEF nested** | GCT GTA AAC ATT TCC TCC AAA TCC |
| **5 prime RACE NP nested** | GGA TCA CTT CCA GCA TTC TTT AG |
| **5 prime RACE P42 nested** | CTC TCC ACT TGT TAG TTG CAT TTC |
| **5 prime RACE NS1 nested** | CCT TCA GCC ACT GCA TCT TC |
| **3 prime RACE SSFP PB2** | CTT GCC TTT GTT GAA GGA TTT CAA G |
| **3 prime RACE SSFP PB1** | CCA AGT TGT TTG CAA CAT GTA CC |
| **3 prime RACE SSFP P3** | GGA AGG ATT CTG TAA CGA GC |
| **3 prime RACE SSFP HEF** | CTC CGA AAA CTG TGA TGC AAG C |
| **3 prime RACE SSFP NP** | GGA AGC GAC GTT CCA AG |
| **3 prime RACE SSFP P42** | GGA AGA CCA AAT GGA AGT TGG G |
| **3 prime RACE SSFP NS1** | GGA GAA GTG GAC TTG TTG TTG C |
| **3 prime RACE PB1 nested** | GTT AAC ACC CCA ATA GGA TCA ATG |
| **3 prime RACE NS1 nested** | CAG ACG ATA CAA ATC TGG ATG TG |

Oligonucleotide sequences for the construction of the reporter fusion cassettes.

| **Name** | **Sequence (from 5' to 3')** |
| --- | --- |
| **pTK Fw** | CGT TGT ACA GGA TCC ATG AAG ACC CTG ATC CTG G |
| **pTK Rv** | GAT CAG CTC GCT CAT GGT GGC GGA TCC TTA AGC GGG |
| **RFP-2A Fw** | ATG AGC GAG CTG ATC AAG GAG AAC ATG CAC |
| **RFP-2A Rv** | GGA TCC TGT ACA ACG CGT CTG CAG CCT AGG |
| **Gaussia Fw** | GCGTTGTACAGGATCCATGGGAGTCAAAGTTCTGTTTGCC |
| **Gaussia Rv** | CCCCAACCCCGGATCTTAGTCACCACCGGCCCC |

Oligonucleotide sequences for cloning of the reporter constructs for minireplicon analysis.

| **Name** | **Primer sequence (From 5' to 3')** |
| --- | --- |
| **pHH21 Amp Fw** | AAT AAC CCG GCG GCC CAA AAT GCC G |
| **pHH21 Amp Rv** | CCC CCC CAA CTT CGG AGG TCG ACC |
| **PB2 NCR GFP Fw** | CCG AAG TTG GGG GGG AGC ATA AGC AGA GGA TGG AGA GCG ACG AGA GCG G |
| **PB2 NCR Gaussia Rv** | GGC CGC CGG GTT ATT AGC AGT AGC AAG AGG ATT TTT TCA ATG TGC TTT AGT CAC CAC CGG CCC CC |
| **PB1 NCR GFP Fw** | CCG AAG TTG GGG GGG AGC ATA AGC AGA GGA TTT TAT AAC AAT GGA GAG CGA CGA GAG CGG |
| **PB1 NCR Gaussia Rv** | GGC CGC CGG GTT ATT AGC AGT AGC AAG AGG ATT TTT CTG TTA TTA AAC AAC GCA AAG CTT AGT CAC CAC CGG CCC CC |
| **P3 NCR GFP Fw** | CCG AAG TTG GGG GGG AGC ATA AGC AGG AGA TTT AAA AAT GGA GAG CGA CGA GAG CGG |
| **P3 NCR Gaussia Rv** | GGC CGC CGG GTT ATT AGC AGT AGC AAG GAG ATT TTT AAC ATT ACA AGG CCT TTG GTT AGT CAC CAC CGG CCC CC |
| **HEF NCR GFP Fw** | CCG AAG TTG GGG GGG AGC ATA AGC AGG AGA TTT TCA AAG ATG GAG AGC GAC GAG AGC GG |
| **HEF NCR Gaussia Rv** | GGC CGC CGG GTT ATT AAT CTT AGA AAA AAT CTC CTT GCT ACT GCT TTA GTC ACC ACC GGC CCC C |
| **NP NCR GFP Fw** | CCG AAG TTG GGG GGG AGC ATA AGC AGG AGA TTA TTA AGC AAT ATG GAG AGC GAC GAG AGC GG |
| **NP NCR Gaussia Rv** | GGC CGC CGG GTT ATT AGC AGT AGC AAG GAG ATT TTT TGT TAA ACA AGA CAA ACC AAC ACC TTT AAC ACC CAC  TGG GGA CTG CAA CAG AAC CAT CCA AAG ATG AGT TAG TCA CCA CCG GCC CCC |
| **P42 NCR GFP Fw** | CCG AAG TTG GGG GGG AGC ATA AGC AGA GGA TAT TTT TGA CGC AAT GGA GAG CGA CGA GAG CGG |
| **P42 NCR Gaussia rv** | GGC CGC CGG GTT ATT AGC AGT AGC AAG AGG ATT TTT TCG CGA TTA GTC ACC ACC GGC CCC C |
| **NS1 NCR GFP Fw** | AGC ATA AGC AGG GGT GTA CAA TTT CAA TAT GGA GAG CGA CG AGA GCG GC |
| **NS1 NCR Gaussia Rv** | AGC AGT AGC AAG GGG TTT TTT TAG TCA CCA CCG GCC CCC TTG |

Oligonucleotide sequences for cloning the polymerase subunits of IDV into the pCAGGS overexpression vector.

| **Name** | **Primer sequence (From 5' to 3')** |
| --- | --- |
| **pCAGGS_RF_Fw** | CTC GAG CTA GCA GAT CTT TTT CCC TCT G |
| **pCAGGS_LF_Rv** | GGT ACC ATG CAT CGA TGA GCT CG |
| **pCAGGS_PB2_Fw** | GCT CAT CGA TGC ATG GTA CCA TGT CAC TAC TAT TAA CG |
| **pCAGGS_PB2_Rv** | AAG ATC TGC TAG CTC GAG TCA AAC TTC CAG ACG |
| **pCAGGS_PB1_Fw** | GCT CAT CGA TGC ATG GTA CCA TGG AAA TAA ACC C |
| **pCAGGS_PB1_Rv (D/660)** | AAG ATC TGC TAG CTC GAG TTA AAC AGA CAG TC |
| **pCAGGS_PB1_Rv (D/CN286)** | AAG ATC TGC TAG CTC GAG TTA AAC AGA CAA TCG |
| **pCAGGS_P3_Fw** | GCT CAT CGA TGC ATG GTA CCA TGT CTA GTG TAA TCA G |
| **pCAGGS_P3_Rv** | AAG ATC TGC TAG CTC GAG TCA TTC AAA GTA C |
| **pCAGGS_NP_Fw** | GCT CAT CGA TGC ATG GTA CCA TGG ACT CAA CAA AAG CCC |
| **pCAGGS_NP_Rv** | AAG ATC TGC TAG CTC GAG TTA ATC TTC ACC |

Oligonucleotide sequences for cloning the viral genomic segments into the pHW2000-vector.

| **Name** | **Primer sequence (From 5' to 3')** |
| --- | --- |
| **pHW2000 Fw** | ACT GCT AAT AAC CCG GCG GCC CAA AAT G |
| **pHW2000 Rv** | GCT TAT GCT CCC CCC CAA CTT CG |
| **IDV segment 1 Fw** | CGG GTT ATT AGC AGT AGC AAG AGG ATT TTT TCA ATG TGC TTC AAA C |
| **IDV segment 1 Rv** | GGG GGA GCA TAA GCA GAG GAT GTC ACT ACT ATT AAC |
| **IDV segment 2 Fw** | CGG GTT ATT AGC AGT AGC AAG AGG ATT TTT CTG TTA TTA AAC AAC GC |
| **IDV segment 2 Rv** | GGG GGA GCA TAA GCA GAG GAT TTT ATA ACA ATG GAA ATA AAC |
| **IDV segment 3 Fw** | CGG GTT ATT AGC AGT AGC AAG GAG ATT TTT AAC ATT ACA AG |
| **IDV segment 3 Rv** | GGG GGA GCA TAA GCA GGA GAT TTA AAA ATG T |
| **IDV segment 4 Fw** | CGG GTT ATT AGC AGT AGC AAG GAG ATT TTT TCT AA |
| **IDV segment 4 Rv** | GGG GGA GCA TAA GCA GGA GAT TTT CAA AGA TG |
| **IDV segment 5 Fw** | CGG GTT ATT AGC AGT AGC AAG GAG ATT TTT TGT TAA ACA AGA CAA ACC AA |
| **IDV segment 5 Rv** | GGG GGA GCA TAA GCA GGA GAT TAT TAA GCA ATA |
| **IDV segment 6 Fw** | CGG GTT ATT AGC AGT AGC AAG AGG ATT TTT TCG |
| **IDV segment 6 Rv** | GGG GGA GCA TAA GCA GAG GAT ATT TTT GAC GC |
| **IDV segment 7 Fw** | CGG GTT ATT AGC AGT AGC AAG GGG TTT TTT C |
| **IDV segment 7 Rv** | GGG GGA GCA TAA GCA GGG GTG TAC AAT TTC AAT |

Oligonucleotide sequences for the full genome sequencing of the rescued virus.

| **Name** | **Primer sequence (From 5' to 3')** |
| --- | --- |
| PB2 938R | GTC TTC GAA CGT TCT TTC TAC TGT TCC |
| PB2 1820R | GAC CGC GTC TTT CAC TTC ATC |
| PB2 1640F | CAC CGA TAG ACA TAG TGG AGA GC |
| PB1 1034R | GTT GGA GCT ACT GAG CAG AGA TC |
| PB1 1674R | GCA GGG CCA TTA AAG CAG TC |
| PB1 1534F | CTC AAA CAT AGC CTT GGA GTT ACC |
| P3 910R | GTT TGC TGC TTC AAA GAA ATC ATA C |
| P3 1627R | GAG TTA ACA CCA CGG CTT CCA C |
| P3 1428F | GAC ACT GAG AGG GGA ATC AAC |
| HEF 1370F | CTT TGA CAC CAG CAA CAA GGA TC |
| HEF 1033R | CTG ATC TTC GTG AAG CTG ACC ACT C |
| HEF 454F | GGG TCA AAC TGG ACT CAG ACA AAG AAG G |
| NP 724F | GGA CCA ATG AGG CAG CTC TGT AAG |
| NP 973R | GTA GCA AGC TGT ATT GAA GGC CAT AAA GG |
| P42 824R | CCA CAG TTT CAG CAT TTA CAC AGG |
| P42 616F | CAC TGC ACA AAG ACG CAG AAG TC |
| NS1 702R | CCA AGC AAC TCC TGA ACC TCT CCA G |
| NS1 64F | CAG AGC AGC AAT CTC CGA ATT GGC |
