## Supplementary material for "Establishment of a reverse genetic system from a bovine derived Influenza D virus isolate": Table S2.docx

**Supplementary Table I:** The nucleotide lengths, percentage of nucleotide homology and number of changed amino acids on each viral segment between the D/bovine/Oklahoma/660/2013 (accession number KF425659-65) and the D/bovine/Switzerland/CN286 virus isolates (Glaus *et al.*, manuscript in preparation).

| Genomic segment | PB2 | PB1 | P3 | HEF | NP | P42 | NS1 |
| --- | --- | --- | --- | --- | --- | --- | --- |
| *Total length in nucleotides* | 2364 | 2330 | 2195 | 2049 | 1775 | 1219 | 868 |
| *Nucleotide homology (in %)* | 97.56 | 98.88 | 98.59 | 95.82 | 98.42 | 96.63 | 98.39 |
| *Amino acid changes* | 4 | 5 | 9 | 25 | 3 | 13 | 5 |
